# Myocardial BDNF regulates cardiac bioenergetics through the transcription factor Yin Yang 1

**DOI:** 10.1101/2021.01.19.427213

**Authors:** Xue Yang, Manling Zhang, Bingxian Xie, Raymond Zimmerman, Qin Wang, An-chi Wei, Zishan Peng, Moustafa Khalifa, Michael Reynolds, Matthew Om, Guangshuo Zhu, Djahida Bedja, Hong Jiang, Michael Jurczak, Sruti Shiva, Iain Scott, Brian O’Rourke, David A. Kass, Nazareno Paolocci, Ning Feng

**Affiliations:** Department of Medicine, Division of Cardiology, Vascular Medicine Institute, University of Pittsburgh, Pittsburgh, PA, USA; Department of Cardiology, Zhongshan Hospital, Fudan University, Shanghai, China; Division of Cardiology, Veteran Affair Pittsburgh Healthcare System, Pittsburgh, PA, USA; Department of Medicine, Division of endocrinology, University of Pittsburgh, Pittsburgh, PA, USA; Echocardiography lab at Heart Center, Ningxia General Hospital, Ningxia Medical University, Ningxia, China; Division of Cardiology, Department of Medicine, Johns Hopkins University, Baltimore, MD, USA; Graduate Institute of Biomedical and Bioinformatics, National Taiwan University, Taiwan

## Abstract

Circulating Brain-derived Neurotrophic Factor (BDNF) is markedly decreased in heart failure patients. Both BDNF and its receptor, Tropomyosin Related Kinase Receptor (TrkB), are expressed in cardiomyocytes, however the role of myocardial BDNF signaling in cardiac pathophysiology is poorly understood. We found that cardiac-specific TrkB knockout (cTrkB KO) mice displayed a blunted adaptive cardiac response to exercise, with attenuated upregulation of transcription factor networks controlling mitochondrial biogenesis/metabolism, including Peroxisome proliferator-activated receptor gamma coactivator 1 alpha (PGC-1α). The cTrkB KO mice developed an exacerbated heart failure progression with transaortic constriction. The downregulation of PGC-1α in cTrkB KO mice exposed to exercise or TAC resulted in decreased cardiac energetics. We further unraveled that BDNF induces PGC-1α upregulation and bioenergetics through a novel signaling pathway, the pleiotropic transcription factor Yin Yang 1 (YY1). Taken together, our findings suggest that myocardial BDNF plays a critical role in regulating cellular energetics in the cardiac stress response.

## Introduction

Endurance exercise has long been linked to cardiovascular health. Multiple recent large-scale prospective or retrospective studies demonstrated that exercise, and the intensity of exercise, is inversely associated with long-term all-cause mortality and CVD mortality ^1–3^. Heart failure is a major public health challenge with high mortality and cost. Exercise training and cardiac rehabilitation have demonstrated numerous benefits for people with heart failure, including improved exercise capacity, increased quality of life, reduced hospitalization, and even decreased mortality rate ^4–6^. Recently launched guidelines for the diagnosis and treatment of acute and chronic heart failure have incorporated a recommendation for regular aerobic exercise in patients with heart failure ^7,8^. However, the molecular mechanism underlying exercise benefits on heart health are still poorly understood, and are pivotal for the development of novel heart failure therapies.

Brain-derived neurotrophic factor (BDNF) is a neurotrophin that regulates energy homeostasis, mitochondrial bioenergetics, and mediates exercise-induced neurogenesis in the brain ^9^. In a prospective study using the Framingham Heart Study cohort, it was found that circulating BDNF levels are inversely associated with the risk of cardiovascular disease ^10^. In addition, retrospective studies show that decreased levels of circulating BDNF are associated with heart failure, worse functional status, higher NT-proBNP, and mortality ^11–13^. BDNF and its receptor, Tropomyosin related kinase receptor B (TrkB), were found in myocardium ^14^, but little is known about the role of myocardial BDNF signaling in cardiac pathophysiology. Our previous study revealed that myocardial BDNF/TrkB signaling is crucial for optimal excitation-contraction coupling ^14^. BDNF has been reported to promote cardiomyocyte survival in myocardial ischemic-reperfusion injury ^15^. Given the emerging evidence that BDNF regulates energy homeostasis in brain and peripheral tissues ^9^, we investigated whether myocardial BDNF/TrkB signaling has any impact on cardiac bioenergetics in response to pathophysiological stress. We found that myocardial BDNF signaling plays a critical role in cardiac bioenergetic regulation in response to exercise or pathological stress, and the upregulation of PGC-1α and other metabolic transcription factors expression appears to be a key underlying mechanism.

## Results

### The cTrkB KO mice display blunted adaptive response to exercise

The chronic exercise-induced cardiac adaptive response protects the heart against pathological stress^16^. The upregulation of Peroxisome proliferator activated receptor gamma coactivator 1 alpha (PGC-1α) expression, the master regulator of metabolic genes and mitochondrial biogenesis, is at the center of adaptive response^16^. PGC-1α coordinates with other metabolic transcription factors, including Peroxisome proliferator-activated receptor α (PPARα) and Estrogen-related receptor α (ERRα), to enhance the fatty acid oxidation by increasing the expression of carnitine palmitoyltransferase 1B (CPT1b), the rate-controlling enzyme of the long-chain fatty acid beta-oxidation pathway in cardiac mitochondria^16^. To investigate if BDNF/TrkB signaling plays a role in the cardiac adaptive response to exercise, myocardial BDNF expression was examined in C57BL/6J mice subjected to swimming exercise. BDNF expression in the heart was significantly increased in C57BL/6J mice subjected to one week of swimming exercise compared to sedentary littermate controls (Fig 1a). To further determine the importance of BDNF/TrkB signaling in cardiac adaptive response, cardiac-specific TrkB KO mice (cTrkB KO mice) were subjected to swimming exercise for one week. We found that while exercise induced increased PGC-1α, PPARα, ERRα, and CPT1b expression in WT mice, the upregulation of these metabolic transcription factors and CPT1b was blunted in cTrkB KO mice (Fig 1b, c). These findings suggested that myocardial BDNF signaling plays a critical role in cardiac adaptive response to endurance exercise.

**Fig 1.**
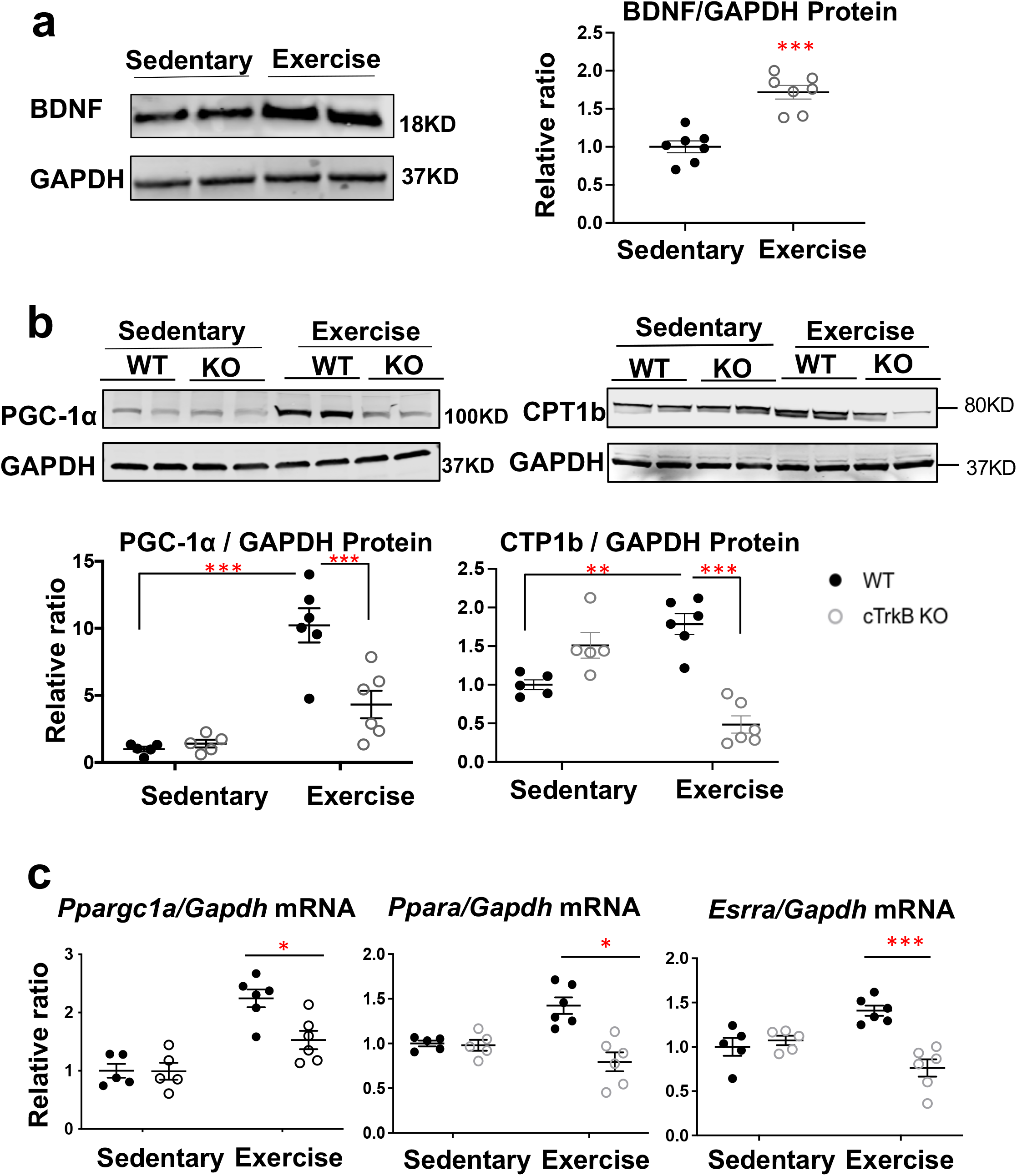
The cTrkB KO mice display blunted adaptive response to exercise. **a.** BDNF expression (n=7) were increased in the mouse hearts with swimming exercise for one week. **b.** The PGC-1α and its downstream target CPT1b protein level were increased with swimming exercise, but the upregulation was blunted in cTrkB KO mice (n=6). **c.** Similarly, the upregulation of PGC-1α mRNA (Ppargc1a), PPARα (Ppara) mRNA, and ERRα (Esrra) mRNA were attenuated in cTrkB KO mice (n=6) subjected to swimming exercise. \**P<0.05,*\*\**P<0.01,*\*\*\**P<0.001*

### Fatty acid oxidation and cardiac function are impaired in cTrkB KO mice subjected to exercise

We next examined whether the blunted PGC-1α and other metabolic transcription factors upregulation in cTrkB KO mice would result in bioenergetics impairment. We measured glucose oxidation and fatty acid oxidation in WT and cTrkB KO mice after two weeks of swimming exercise, using isolated working heart system with C^14^-labeled glucose and H^3^-labeled oleic acid as described previously^17^. We found that fatty acid oxidation was impaired in cTrkB KO mice, compared to WT controls (Fig 2a). There was no significant difference in glucose oxidation (Fig 2a). In consistent with the decreased fatty acid oxidation in cTrkB KO mice, the contractility of cTrkB KO mice hearts, measured by pressure catheter, was decreased too (Fig 2b). We also measured cardiac function in cTrkB KO mice after swimming exercise by echocardiography. Our previous study showed there was no difference in cardiac function at baseline^14^. We further followed up cardiac function of cTrkB KO mice until one year old. The LV systolic function of cTrkB KO mice was preserved at one-year old, and there is no difference in cardiac function between WT littermate controls and cTrkB KO mice (Fig S1). Echocardiography study demonstrated the cardiac function in cTrkB KO mice was decreased after 2 weeks of swimming exercise, while the cardiac function in WT controls was preserved (Fig 2c). Taken together, our findings highlighted that myocardial BDNF/TrkB signaling is required for the adaptive response to endurance exercise.

**Fig 2.**
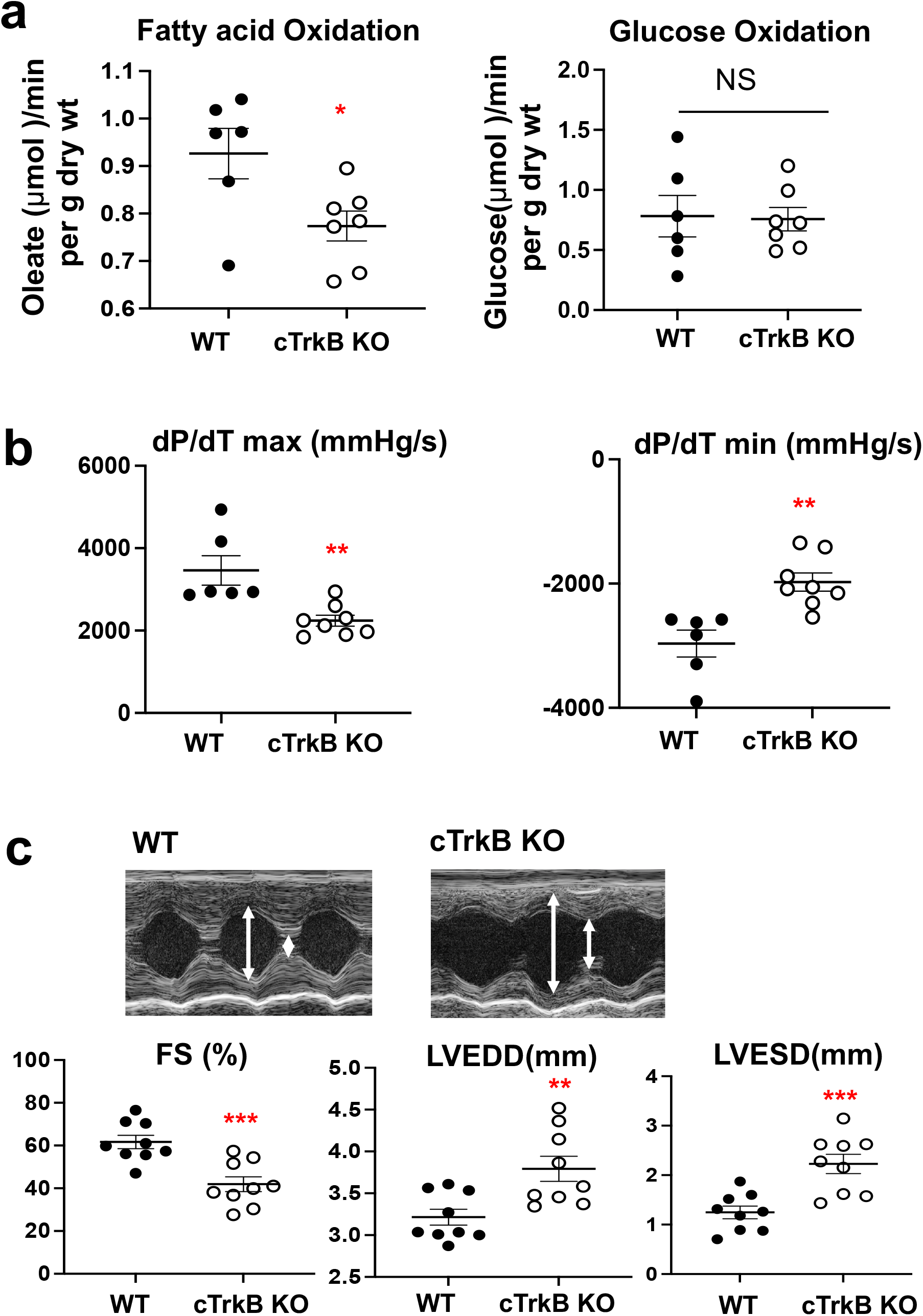
The Fatty acid oxidation rate and cardiac function are impaired in cTrkB KO mice with swimming exercise. **a.** Compared to WT controls, the fatty acid oxidation rate, measured in isolated working heart with H^3^-labeled oleic acid, was decreased in cTrkB KO mice with 2 weeks of swimming exercise, whereas there was no significant change in glucose oxidation (n=6-7). **b.** Compared to WT controls, the contractility of hearts from cTrkB KO mice with exercise, which was measured by dP/dT in the isolated working heart system, was significantly reduced (n=6-7). **c.** Representative echocardiography and data summary of WT and cTrkB KO mice subjected to swimming exercise. Compared to WT mice, cTrkB KO mice showed decreased fractional shortening, increased LV end systolic or diastolic dimension (n=9). \**P<0.05,*\*\**P<0.01,*\*\*\**P<0.001*

### The cTrkB KO mice show exacerbated heart failure progression against pressure overload

To inspect whether BDNF signaling plays a role in heart failure progression, we examined cardiac BDNF expression in C57BL6J mice subjected to transaortic constriction (TAC). We found that although there were no significant changes in the expression of BDNF at the concentric hypertrophy stage (2 weeks after TAC), the BDNF level was significantly decreased at the heart failure stage (6 weeks after TAC) (Fig 3a). Consistently, we found that myocardial BDNF level was decreased in explanted hearts from patients with non-ischemic cardiomyopathy Fig 3a), suggesting impaired BDNF signaling may be an important mechanism for heart failure progression. Next, to determine the importance of BDNF signaling in heart failure progression, both male and female cTrkB KO mice were subjected to TAC. Compared to WT littermate controls as well as age matched MHC-Cre mice, male cTrkB KO mice began to display worse cardiac function at 10 days post TAC. Cardiac function in male cTrkB KO mice further deteriorated at 6 weeks post-TAC with decreased fractional shortening (WT 43.1±2.8% [n=12] or MHC-Cre 38.7±5.2% [n=9] vs cTrkB KO 23.1±2.1% [n=15], *P< 0.001 and P<0.01 respectively*), (Fig 3b, c) and increased dilation of left ventricular chamber (Fig 3b, c). Because there was no difference in cardiac function measured by echocardiography between WT littermate controls and MHC-Cre controls, we combined heart/lung weight data, as well as fetal gene expression together as one group. Compared to WT controls, the heart weight and lung weight were increased (Fig 3d), and the fetal gene reprogramming was augmented in TAC cTrkB KO mice (Fig 3e). Consistent with this, female cTrkB KO mice showed accelerated heart failure progression after TAC compared to WT controls (Fig S2). Our findings showed that myocardial BDNF signaling is essential to protect the heart against pathological stress.

**Fig 3.**
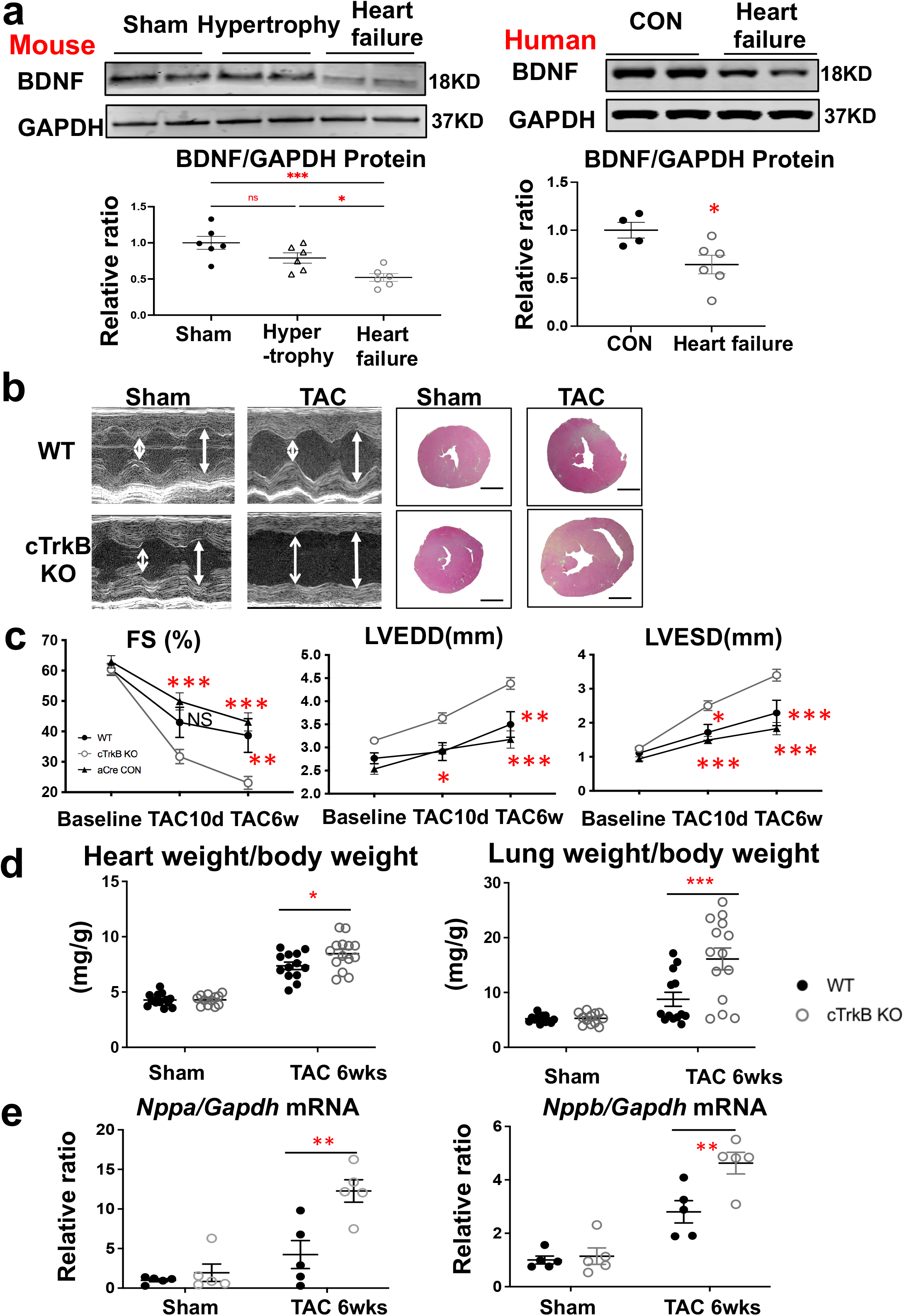
cTrkB KO mice display exacerbated heart failure progression post TAC. **a.** In contrast with exercise, there was no significant change in myocardial BDNF level in concentric cardiac hypertrophy (2 weeks after TAC), and the BDNF was significantly decreased in failing hearts (6 weeks after TAC) (n=6). The level of myocardial BDNF was also significantly reduced in patients with non-ischemic cardiomyopathy (n=4-6) **b.** Representative echocardiography and histology of WT and cTrkB KO mice (scale bar 1mm) subjected to sham or TAC surgery. **c.** Compared to WT mice and MHC-Cre controls, cTrkB KO mice showed decreased fractional shortening (WT 43.1±2.8% or MHC-Cre 38.7±5.2% vs cTrkB KO 23.1±2.1%), increased LV end diastolic dimension (LVEDD) and LV end systolic dimension (LVESD). (WT n=12, MHC-Cre n=9, cTrkB KO n=15). **d.** TAC cTrkB KO mice heart weight and lung weight were increased relative to WT controls. (WT n=12, cTrkB KO n=9) **e.** ANP (Nppa) and BNP (Nppb) were increased in cTrkB KO mice subjected to TAC (n=6). \**P<0.05,*\*\**P<0.01,*\*\*\**P<0.001*

### PGC-1α expression, mitochondrial proteins level, DNA copy number and function are significantly reduced in cTrkB KO mice subjected to TAC

Given that cardiac BDNF regulates PGC-1α and energetics in response to exercise, we next investigated whether the regulation of PGC-1α by BDNF is also critical in the cardiac response to pathological stress. In our experimental model, we observed that PGC-1α expression (mRNA and protein expression) was not changed in WT mice after TAC (Fig 4a, b). We found PGC-1α, along with other metabolic transcription factors such as PPARα, ERRα, and Transcription factor A, mitochondrial (TFAM), were significantly decreased in TAC cTrkB KO mice hearts compared to WT controls, suggesting myocardial BDNF is essential in regulating cardiac metabolism in response to pathological stress (Fig 4a, b and Fig S3).

**Fig 4.**
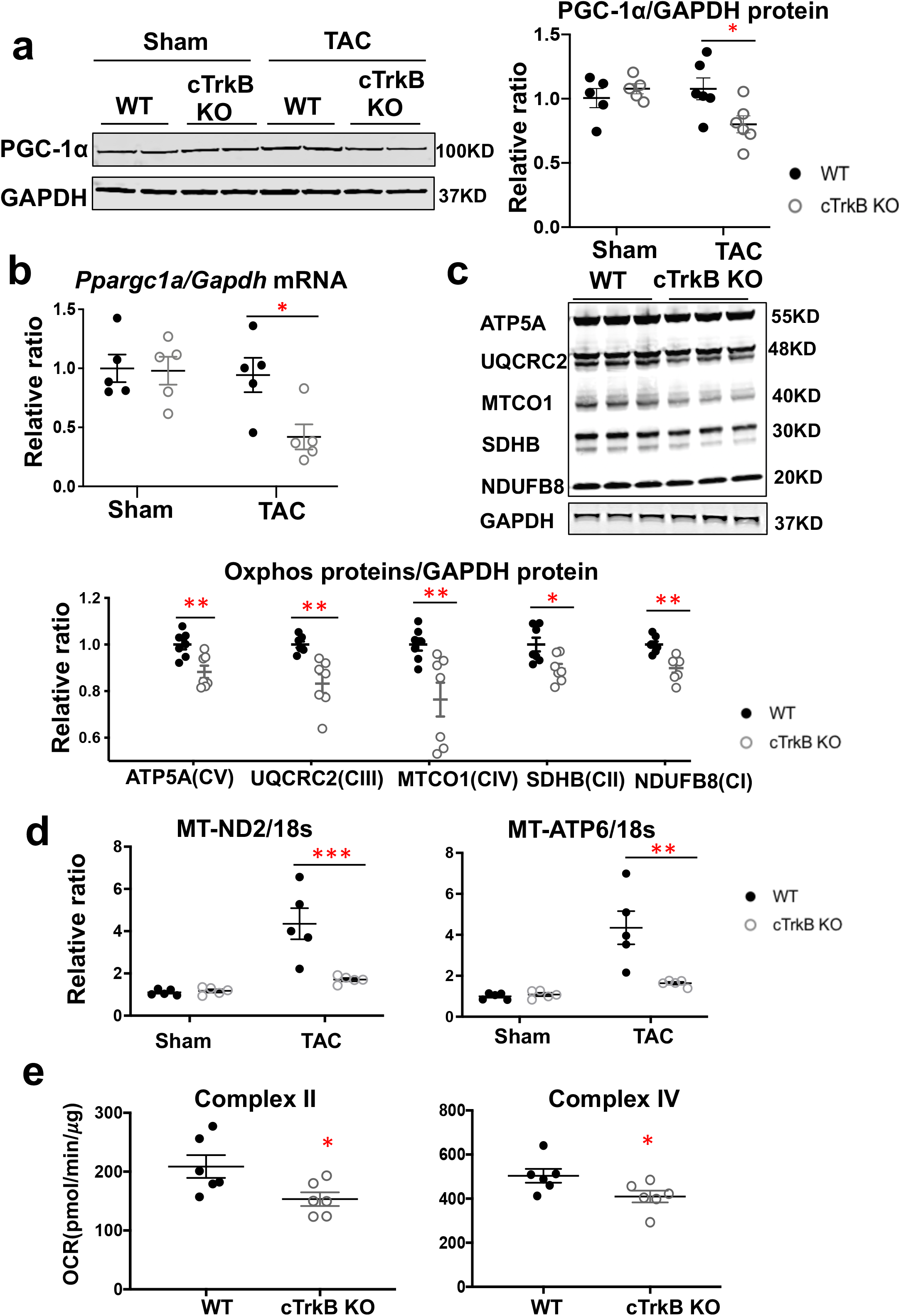
PGC-1α expression, Mitochondrial proteins, mitoDNA copy numbers, and oxygen consumption rate are decreased in cTrkB KO mice subjected to TAC. **a., b.** The PGC-1α mRNA (Ppargc1a) and protein level were significantly lower in cTrkB KO mice subjected to TAC compared to WT controls (n=5-6). **c.** OXPHOS cocktail western blot showed that mitochondrial proteins including NDUFB8 (complex I), SDHB (complex II), UQCRC2 (complex III), ATP5A (complex IV), and COXII (complex V) were decreased in TAC cTrkB KO mice, compared to WT controls (n=6). **d.** Mitochondrial DNA copy number, assessed by MT-ND2/18s RNA gene and MT-ATP6/18s RNA gene, was reduced in cTrkB KO mice subjected to TAC (WT n=5, cTrkB n=5). **e.** oxygen consumption rate of complex II and complex IV were impaired in TAC cTrkB KO hearts relative to WT controls. (n=6) \**P<0.05,*\*\**P<0.01,*\*\*\**P<0.001*

We next examined whether downregulation of PGC-1α and the other nuclear transcription factors result in reduced mitochondrial biogenesis/content, and impaired mitochondrial respiratory function in cTrkB KO mice. Mitochondrial protein abundance was assessed by Oxphos Western Blot cocktail antibodies. ATP Synthase F1 Subunit Epsilon (ATP5A), Ubiquinol-Cytochrome C Reductase Core Protein 2 (UQCRC2), mitochondrially-encoded Cytochrome C Oxidase I (MTCO1), Succinate Dehydrogenase Complex Iron Sulfur Subunit B (SDHB), and NADH:Ubiquinone Oxidoreductase Subunit B8 (NDUFB8) were all significantly decreased in cTrkB KO mice subjected to TAC (Fig 4c). Mitochondrial biogenesis was assessed by mitochondrial DNA (Mitochondrially Encoded NADH:Ubiquinone Oxidoreductase Core Subunit 2, MT-ND2, or Mitochondrially Encoded ATP Synthase Membrane Subunit 6, MT-ATP6) to genomic DNA (18s RNA) ratio^18^. In our experimental model, mitoDNA copy number was increased in TAC WT mice, compared to sham WT mice. (Fig 4d). Relative to WT controls, mitochondrial DNA copy number was markedly decreased in stressed cTrkB KO mice hearts (Fig 4D). In addition, we evaluated mitochondrial respiratory function on freshly isolated mitochondria in TAC cTrkB KO mice using the Seahorse XF Bioanalyzer as described previously^19^. Complex II and complex IV oxygen consumption rate (OCR) were significantly decreased in cTrkB KO mice subjected to TAC (Fig 4e), while Complex I OCR was preserved (Fig S4). Collectively, consistent with markedly downregulated energy-related nuclear transcription factors, mitochondrial function, content, and biogenesis were impaired in cTrkB KO mice subjected to TAC.

### BDNF/TrkB signaling regulates PGC-1α expression through AKT-mTOR-YY1 pathway

To further demonstrate that PGC-1α is the direct target of BDNF/TrkB signaling, BDNF was overexpressed with plasmid or TrkB was knocked down with siRNA in neonatal rat cardiomyocytes (NRCMs). The efficiency of knockdown was validated by RT-PCR (Fig S5). We found that β-adrenergic stimulation with isoproterenol increased BDNF expression (Fig 5a). Consistent with the *in vivo* studies, we found TrkB knockdown attenuated isoproterenol induced PGC-1α upregulation (Fig 5b). Overexpression of BDNF increased PGC-1α expression (Fig 5c).

**Fig 5.**
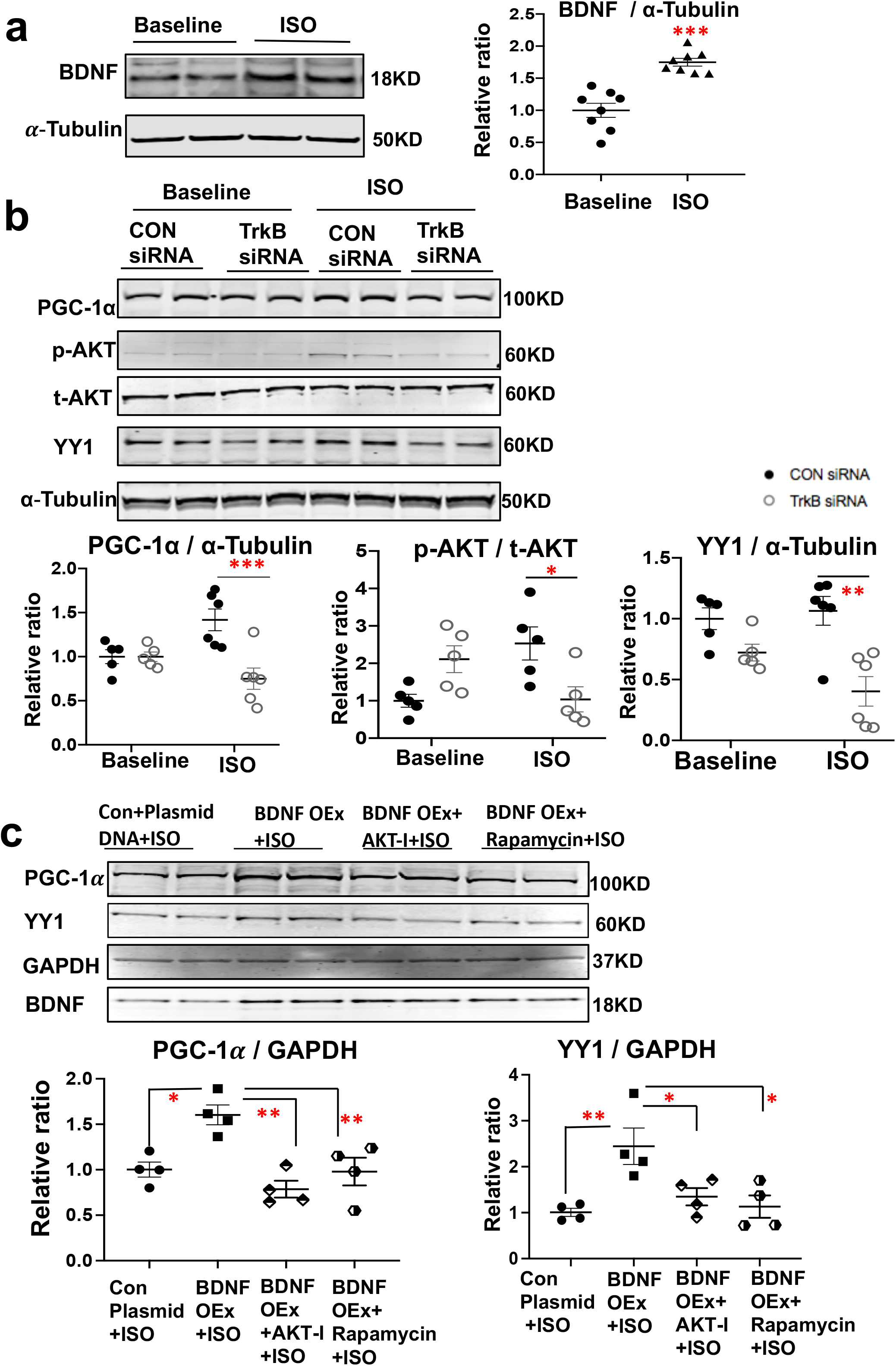
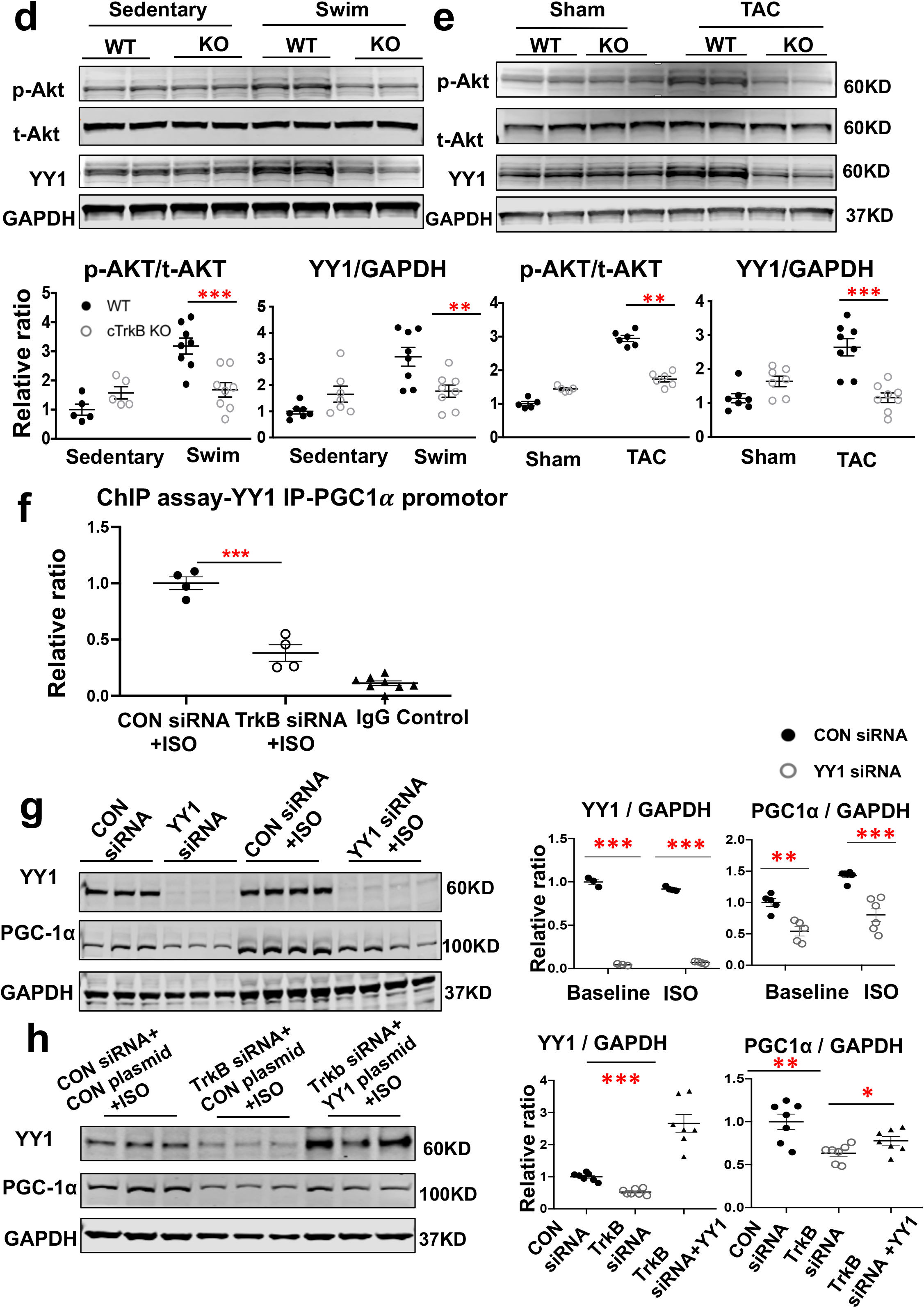
Myocardial BDNF regulates PGC-1α expression through AKT-mTOR-YY1 pathway. **a.** Isoproterenol (10μM) induced BDNF expression in NRCMs (n=8). **b.** PGC-1α and transcription factor Yin-yang 1 (YY1) expression, and the level of phosphorylated AKT were increased with isoproterenol stimulation, and the upregulation was attenuated in TrkB siRNA transfected NRCMs (n=5-6). **c.** Overexpression of BDNF increased PGC-1α and YY1 expression, and the BDNF induced upregulation of PGC-1α and YY1 were attenuated by AKT inhibitor LY294002 or mTOR inhibitor rapamycin (n=4). **d and e.** The level of phosphorylated AKT and YY1 expression were elevated in WT mice responding to swimming exercise or TAC (n=5), but the elevation was blunted in cTrkB KO mice (n=6-8). **f.** Chromatin Immunoprecipitation Assay demonstrated the binding of YY1 to PGC-1 promoter was significantly decreased in NRCMs with TrkB knockdown with the incubation of isoproterenol (n=4). **g and h.** YY1 knockdown decreased PGC-1α expression in NRCMs with or without isoproterenol (n=4-6), and the overexpression of YY1 in NRCMs recovered the downregulation of PGC-1α expression in NRCMs with TrkB knockdown (n=7). *P<0.05,*\*\**P<0.01,*\*\*\**P<0.001*

We investigated the signaling pathways mediating BDNF-induced PGC-1α expression. It has been reported that Erk/CREB mediates BDNF-induced PGC-1α upregulation in the brain^20^. However, there was little change in CREB activation in cTrkB KO mice subjected to exercise or TAC, and the p-Erk level was increased in TAC cTrkB KO mice (Fig S6a). In addition, chromatin immunoprecipitation assay (ChIP) showed there was no difference in the binding of CREB to PGC-1α promotor between WT and cTrkB KO mice subjected to TAC (Fig S6b). AMPK, an important regulator of PGC-1α, is implicated in BDNF induced fatty acid oxidation enhancement in skeletal muscle responding to contraction ^21^. We therefore investigated whether AMPK is involved in BDNF-induced PGC-1α upregulation in the heart, but found that there was little change of phosphorylated or total AMPK level in cTrkB KO mice subjected to exercise or TAC (Fig S6a). Instead, we found that phosphorylated AKT (p-AKT, Thr308) was markedly decreased in cTrkB KO mice subjected to exercise or TAC, without significant change in total AKT level (t-AKT) (Fig 5d and 5e). Consistently, TrkB knockdown in NRCMs attenuated isoproterenol-induced AKT phosphorylation (Fig 5b). Importantly, AKT inhibition with LY294002 prevented BDNF overexpression induced PGC-1α upregulation (Fig 5c). AKT/mTOR is a common downstream signaling pathway of growth factor/tyrosine kinase receptors, and it was reported that Mechanistic Target of Rapamycin Kinase (mTOR) coordinates cellular growth and energy metabolism by controlling PGC-1α and other metabolic gene transcription ^22^. We therefore investigated whether mTOR is involved in BDNF-induced PGC-1α expression. Indeed, mTOR inhibition with rapamycin blunted BDNF overexpression induced PGC-1α (Fig 5c). The mTOR regulates PGC-1α expression by promoting Yin Yang transcription factor 1 (YY1) binding to the PGC-1α promoter ^22^. We found that YY1 was significantly increased in WT mice subjected to exercise or TAC, however, the upregulation of YY1 was blunted in cTrkB KO mice (Fig 5d and 5e). TrkB knockdown in NRCMs attenuated isoproterenol-induced YY1 upregulation (Fig 5b). BDNF overexpression in NRCMs increased YY1 expression, and AKT inhibition with LY294002 or rapamycin blunted BDNF overexpression induced YY1 upregulation (Fig 5c). Moreover, we showed the binding of YY1 to PGC-1α promoter was decreased in NRCMs with TrkB knockdown using chromatin immunoprecipitation assay (ChIP, Fig 5f). Taken together, these findings suggested BDNF controls PGC-1α transcription through AKT-mTOR-YY1 pathway.

Because YY1 could be either transcription activator or repressor^23^, we examined whether YY1 promotes PGC-1α transcription in cardiac myocytes. Indeed, YY1 knockdown decreased PGC-1α expression in NRCMs with or without isoproterenol stimulation (Fig 5g). Importantly, overexpression of YY1 at least partially recovered the downregulation of PGC-1α expression in NRCMs with TrkB knockdown, demonstrating YY1 is the key mediator of BDNF regulation on PGC-1α expression (Fig 5h). Therefore, our study provided a new downstream signaling pathway of BDNF signaling.

### TrkB or YY1 knockdown in NRCMs impairs mitochondrial function

To investigate whether there is a functional consequence of decreased PGC-1α expression in the NRCMs with TrkB knockdown, we examined the mitochondrial function in NRCMs with TrkB knockdown using Seahorse XF system (Seahorse Bioscience). Without isoproterenol stimulation, there was no difference in oxidative respiratory capacity in NRCMs with TrkB knockdown, but the oxygen consumption rate was significantly decreased in NRCMs with TrkB knockdown at basal respiratory condition in the presence of isoproterenol (Fig 6a). These findings further demonstrated BDNF/TrkB signaling regulates cardiomyocytes PGC-1α expression and energetics. Next, to determine whether there is a functional consequence of decreased PGC-1 expression in the NRCMs with YY1 knockdown, we inspected the mitochondrial function in NRCMs with YY1 knockdown. With YY1 knockdown, the oxidative respiratory capacity was markedly decreased under both basal condition and maximum capacity, with and without isoproterenol stimulation (Fig 6b), demonstrating YY1 plays an essential role in regulating bioenergetics in cardiac myocytes.

**Fig 6.**
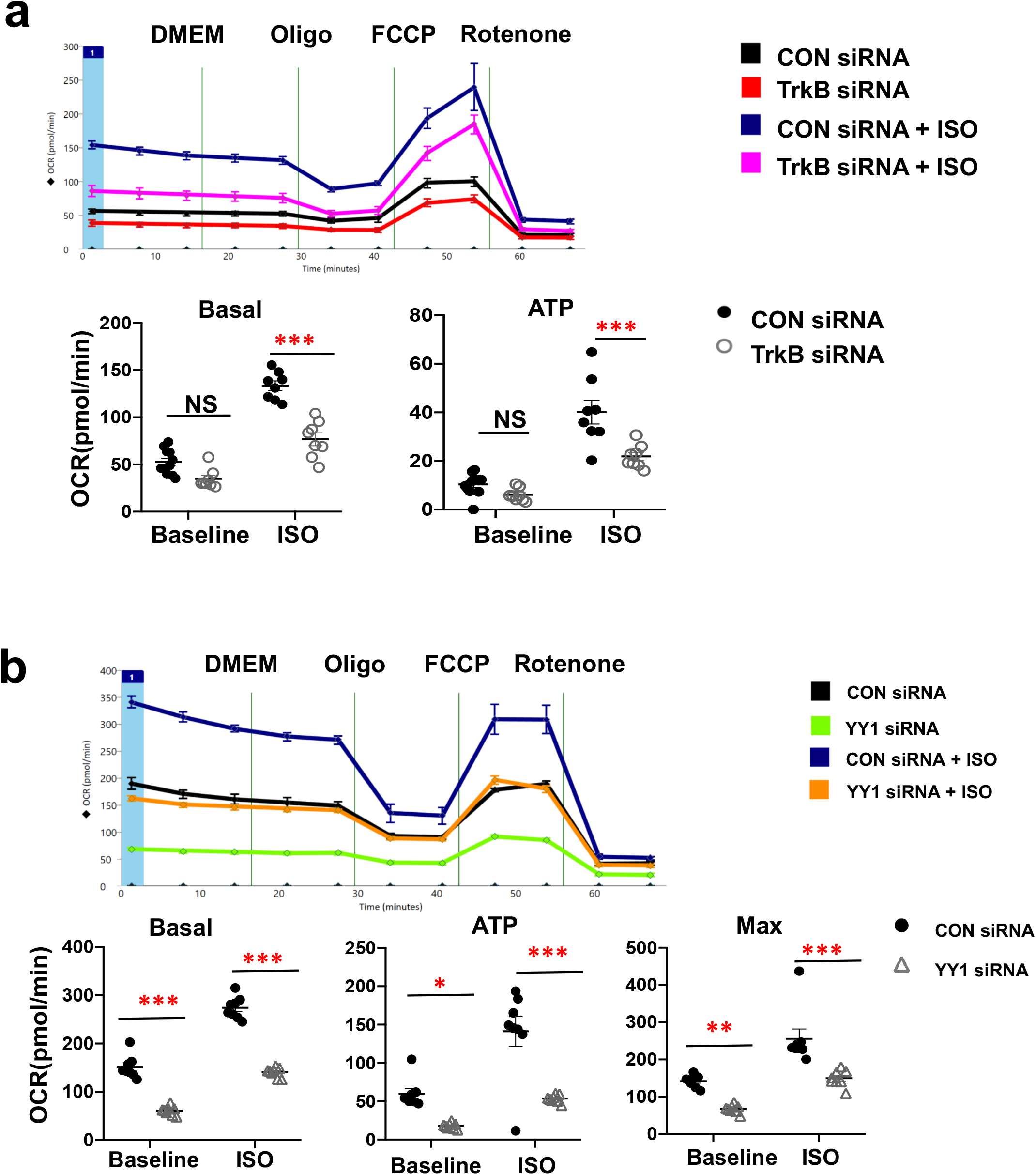
TrkB or YY1 knockdown in NRCMs impairs mitochondrial function. **a.** While there is no significant difference in oxidative respiratory capacity without isoproterenol stimulation between control and NRCMs with TrkB knockdown, the basal respiratory function was significantly decreased with TrkB knockdown in the presence of isoproterenol (n=8). **b.** NRCMs transfected with YY1 siRNA displayed both impaired basal and maximum respiration, at both baseline and with isoproterenol incubation (n=8). \**P<0.05,*\*\**P<0.01,*\*\*\**P<0.001*

## Discussion

The role of myocardial BDNF signaling in cardiac pathophysiology is poorly understood. Here, we showed that myocardial BDNF signaling was enhanced in response to endurance exercise but was impaired in failing hearts. Using cardiac-specific TrkB KO model, we demonstrated that myocardial BDNF signaling played an essential role in the adaptive response to exercise, and protected against heart failure progression induced by chronic hemodynamic stress. Our findings suggested myocardial BDNF signaling controlled the abundance of cardiac metabolic transcription regulators including PGC-1α, PPARα, and ERRα in response to exercise and pathological stress. These findings suggest that myocardial BDNF signaling-elicited bioenergetics enhancement may be one of the underlying molecular mechanisms in the exercise-induced adaptive response and exercise-granted cardiovascular protection. Developing therapeutic drugs targeting the BDNF signaling pathway may provide exercise-mimic beneficial effects against pathological stress, especially for patients with limited exercise capacity. The therapeutic approach targeting BDNF signaling, thereby targeting cardiac metabolism/bioenergetics, differs from standard heart failure therapy aiming at neurohormone inhibition, and therefore has the potential to provide additional or synergistic benefit for heart failure patients who are already on standard therapies. Because impaired BDNF signaling plays a critical role in many neurodegerative diseases, there has long been a major interest to develop TrkB agonists. Currently there are a number of small molecular compounds or peptide-based TrkB agonists proven effective in animal models of neurodegenerative diseases ^24–27^. Our findings suggest these TrkB agonists could be potentially repurposed for heart failure therapy.

We observed that BDNF expression is increased in the heart in response to exercise. However, which cell type is responsible for this enhanced BDNF level in heart remains unclear, since BDNF is expressed in nerves ^15^, vascular endothelial cells ^28^ and vascular smooth muscle cells, and cardiac myocytes ^14,15^. Our cultured cardiac myocytes studies suggested paracrine or autocrine of BDNF in the cardiomyocytes may at least play some role. Because we were not certain that cardiomyocyte-derived BDNF is the major resource of BDNF in the heart, cTrkB KO mice were selected to investigate the importance of BDNF signaling in cardiac physiology. However, TrkB also interacts with other neurotrophins including neurotrophin-3 and neurotrophin 4/5. Therefore, it is quite possible that other neurotrophins contribute to activate TrkB receptor in response to cardiac stress in addition to BDNF.

In contrast to exercise, we found chronic hemodynamic stress results in decreased BDNF expression in the late stage of heart failure. The molecular mechanism of this decreased BDNF expression is unclear. BDNF expression is also decreased in neurodegenerative diseases^29^, and two mechanisms are often involved in this impairment. One mechanism is increased methylation of the BDNF promotor, which leads to enhanced binding of MeCP2, a transcriptional suppressor, to the BDNF promotor ^30^. Another mechanism of BDNF downregulation is increased miR-212/132 binding to 3’ UTR of BDNF mRNA ^31^. Enhanced MeCP2 and miR-212/132 signaling have been reported to play an important role in cardiac hypertrophy ^32,33^. Whether these two mechanisms of BDNF suppression are also involved in the failing heart needs to be investigated.

Our study provided one new downstream signaling pathway mediates BDNF action. We found that cardiac BDNF signaling activated AKT/mTOR, promoting the upregulation of YY1 expression in response to pathophysiological stress. YY1 expression in the heart was increased with exercise or TAC, but the upregulation was attenuated in cTrkB KO mice. Consistently, in cultured cardiomyocytes, knockdown of TrkB attenuated β-adrenergic stimulation induced YY1 and PGC-1α upregulation, and the binding of YY1 to PGC-1α promoter. Importantly, YY1 overexpression rescued TrkB knockdown induced PGC-1α downregulation, demonstrating YY1 is the key mediator of BDNF induced PGC-1α expression. YY1 plays a critical role in cardiac development through a number of mechanisms^34,35^. However, the role of YY1 in adult cardiac physiology is poorly understood. YY1 has been shown to attenuate phenylephrine-induced cardiomyocytes hypertrophy by preventing histone deacetylase 5 (HDAC5) phosphorylation and subsequent exporting out of nuclear^36^. HDAC5 acts as a repressor of the cardiac hypertrophy through inhibition of myocyte enhancer factor 2 (MEF2)^37^. Here we showed that YY1 is implicated in cardiac energetics enhancement in response to exercise and pressure overload. Indeed, knocking down of YY1 in cardiomyocytes resulted in blunted PGC-1α upregulation and mitochondrial respiration enhancement by β-adrenergic stimulation. Therefore, our study revealed a new role of YY1 in the heart and suggested that the YY1 activation could be used to screen TrkB agonists for heart failure therapy.

Interestingly, we observed that the level of p-AKT in the heart were markedly increased in both WT mice subjected to exercise and TAC, which is consistent with previous studies showing that AKT pathway is critical in both adaptive and maladaptive hypertrophy^38^. In response to exercise, AKT is activated by growth factors, such as insulin-like growth factor 1 (IGF-1), and the exercise induced AKT activation promotes cardiac metabolism and adaptive cardiac hypertrophy^38^. In response to pathological stress, the heterotrimeric G-proteins Gq and G11 signalings are activated by prohypertrophic G protein receptor agonists including angiotensin II, endothelin or norepinephrine. Gq or G11 activation can also turn on AKT signaling to transduce maladaptive hypertrophy. The Gq or G11 derived AKT activation does not increase PGC-1α expression or enhance cardiac energetics^38^. Our findings on the cardiac protection of BDNF/TrkB elicited AKT activation adds evidence to support that one major determinant of the downstream targets as well as the adaptive versus maladaptive growth of AKT signaling is likely to be the upstream input. Similarly, we found that the myocardial YY1 expression were upregulated responding to exercise and TAC, whereas PGC-1α expression was significantly upregulated in hearts with exercise but had little change or was deceased in TAC hearts. YY1 is a dual functional transcription factor that can act as a transcriptional repressor, a transcriptional activator, or a transcriptional initiator^39^, depending on the context. The discrepancy of YY1/PGC-1α expression in the hearts between exercise and TAC further supports that the transcriptional activator versus repressor of YY1 is dependent on the upstream input and possibly the recruitment of additional signaling by the upstream input. We speculate that the BDNF/TrkB upregulates PGC-1α transcription not only through the increase of YY1 expression and binding to PGC-1α promoter, but also through additional cofactors recruitment to the PGC-1α promoter.

Recently it has been found that NTRK gene fusions including either NTRK1, NTRK2 or NTRK3 (encoding the neurotrophin receptors TrkA, TrkB and TrkC, respectively) are oncogenic drivers of various adult and pediatric tumour types, and Trk inhibitors were developed to treat NTRK-fusion positive cancers ^40^. Our findings suggest that these inhibitors need to be used cautiously in the patients with cardiac dysfunction.

In summary, our work revealed a novel role of cardiac BDNF signaling, and further investigation may yield several clinical applications.

## Methods

### Animal models and procedures

Study procedures were approved by the Johns Hopkins University and University of Pittsburgh Animal Care and Use Committees in accordance with National Institutes of Health guidelines. Floxed TrkB mice were generated as reported previously ^41^ and were kindly provide by Dr. David Ginty at Harvard University. Cardiac-specific TrkB KO mice (cTrkB KO mice) were generated by crossing with αMHC-Cre mice (Jackson Lab, Stock # 011038) as we reported previously^14^.

WT littermate controls and cTrkB KO mice were subjected to swimming exercise in 32 °C water, 90 minutes per session, twice a day for one week (Gene expression studies) or two weeks (Fatty acid and Glucose oxidation studies), after one week of training session. TAC procedure was performed as previously described ^42^. Briefly, Sterile surgery was performed in our dedicated facility. Anesthesia was induced with 3.0% isoflurane, and maintained by ventilator at 2.0% isoflurane. Skin was shaved and cleaned with povidone-iodine. The chest is entered by a limited incision immediate left to the sternum to expose the mid-thoracic aorta. A 7-0 proline suture tie was placed around the aortic arch between bronchiocephalic artery and left common carotid artery and tightened around a 27 or 26 gauge needle - a common diameter for aortic banding in the mouse. Once the tie was fixed, the needle was removed, and the suture/stenosis remained. The chest was then closed with 6-0 proline suture and 5-0 silk suture. Serial echocardiography were performed in conscious mice (VisualSonics, Vevo 3100) as described previously^43^.

### Isolated working heart study

The isolated working heart study was performed as previously described with modification^17^. Hearts from anesthetized mice were rapidly excised and cannulated via the aorta and pulmonary vein/left atrium in warm oxygenated Krebs-Henseleit buffer (118 mM NaCl, 25 mM NaHCO3, 0.5 mM Na-EDTA [Disodium salt dihydrate], 5 mM KCl, 1.2 mM KH2PO4, 1.2 mM MgSO4, 2.5 mM CaCl2, 11 mM glucose), and anterograde perfusion was initiated with a constant atrial pressure of 11 mmHg against an aortic workload of 50 mmHg. Left ventricle pressure was measured via Mikro-tip pressure catheter (Millar) carefully inserted into the LV through the aorta. Hearts were paced at a rate slightly above intrinsic rate (~360-500 bpm). The work-performing heart was perfused with Krebs-Henseleit buffer with the supply of ^14^C-radiolabeled glucose ([U-^14^C]Glucose) and a ^3^H-radiolabeled oleic acid ([9,10-^3^H]oleic acid). After 30 minutes, the coronary effluent was collected, and the flow rate was recorded. The end byproduct ^14^CO2 and ^3^H2O were quantitatively recovered from the coronary effluent for the final calculation. The left ventricular pressure was measured and calculated as ±dP/dT throughout the experiments.

### Mitochondrial isolation and mitochondrial physiology study

Mitochondria from mouse hearts were isolated as described previously ^19^. Briefly, mouse hearts were homogenized in isolation solution 75 mM sucrose, 225 mM mannitol, and 1 mM EGTA, pH 7.4, with 0.1mg/ml proteinase. The proteinase was neutralized with 3 volume of isolation solution containing 0.2% fatty acid–free BSA (Sigma-Aldrich). After a first centrifugation (500 *g* for 10 min) to discard unbroken tissue and debris, the supernatant was centrifuged at 10,000 *g* for 10 min to sediment the mitochondria, which were washed twice thereafter by centrifugation at 7,700 *g* for 5 min each, the first one with isolation solution in presence of BSA, and the second in absence of BSA.

In brief, mitochondria were assayed in polyethyleneimine-coated XF96 plates using the Seahorse XF system (Seahorse Bioscience) as previosly described ^44^. After removing the solution of polyethyleneimine, the equivalent of 5–15 μg mitochondrial protein was transferred to each well. The status of the different complexes of the respiratory chain was evaluated with substrates of Complex I (5 mmol/L glutamate/malate), Complex II (5 mmol/L succinate and 1μmol/L rotenone) and Complex IV (0.5 mmol/L N,N,N,N-tetramethyl-p-phenylenediamine [TMPD] and 3 mmol/L sodium ascorbate). Maximal respiratory capacity was analyzed in the presence of 1 mmol/L ADP and 50 μmol/L dinitrophenol.

### Mitochondrial DNA copy number and mitochondrial protein level assessment

Mitochondrial DNA and genomic DNA were isolated from mouse hearts using DNeasy Blood & Tissue Kits (Qiagen) according to the manufacturer’s instructions. MitoDNA copy number was assessed with Mitochondrially Encoded NADH:Ubiquinone Oxidoreductase Core Subunit 2, (MT-ND2), or Mitochondrially Encoded ATP Synthase Membrane Subunit 6, (MT-ATP6)/genomic DNA (18s RNA genes) ratio as described^18^. The Taqman primers were obtained from Thermofisher Scientifics (MT-ATP6: Mm03649417_g1, MT-ND2: Mm04225288_s1, 18 sRNA gene: Hs99999901_s1). Mitochondrial proteins level was evaluated by Western Blot using OXPHOS cocktail antibodies (Abcam, #ab110413).

### Cardiac myocytes studies

Neonatal rat cardiac myocytes (NRCMs) were freshly isolated as described previously^43^. Briefly, the hearts were quickly removed from one to three days old Sprague Dawley neonates and immersed into chilled KH buffer (pH 7.5) containing: NaCl 140mM, KC1 4.8mM, MgSO4-7H2O 1.2mM, dextrose 12.5mM, NaHCO3 4mM, NaH2PO4 1.2mM, and HEPES 10mM. The ventricles were cut into 1-2mm pieces and the cardiac myocytes isolation was achieved by digestion with 0.04% trypsin and type II collagenase (Worthington #LS004176) 0.4mg/ml in KH buffer at 37°C. The digestion was terminated by DMEM containing 10% fetal bovine serum (FBS). Non-cardiomyocyte cells were removed by rapid attachment (90 minutes incubation in culture dishes). Cardiomyocytes were plated at the density of 2X10^5^/ml in DMEM containing 10%FBS and 0.1mM BrdU to prevent the growth of fibroblasts. 24 hours after cells plating, the medium was changed to serum free DMEM containing 0.1% Insulin-transferrin-selenium (Thermo Fisher Scientifics).

### Mitochondrial bioenergetics measurements in cutured cardiac myocytes

Oxygen consumption rate (OCR) was measured in NRCMs transfected with control siRNA, TrkB siRNA or YY1 siRNA using the Seahorse XF system (Seahorse Bioscience) as described previously^45^. Basal OCR in each well was measured, followed by serial treatment with FCCP (7.5 μmol/L) and rotenone (2 μmol/L). Each experiment was repeated to ensure reproducibility, and the data presented are technical replicates (N = 8) of a single representative study.

### Western blot

Protein extracts were prepared in RIPA lysis buffer (Thermo Fisher Scientific) from snap-frozen heart tissues. Protein concentration was measured by BCA assay (Thermo Fisher Scientifics). The protein electrophoresis were performed on 4–12% Bis-Tris NuPage gels (Thermo Fisher Scientifics) or Bio-Rad Mini-PROTEAN® TGX™ Precast Gels. The Bio-Rad Trans-Blot Turbo Transfer System was used for the proteins transfer to nitrocellulose membranes. Secondary antibodies used were from LI-COR Biosciences. The blots were quantified using Image J or LI-COR Image studio Lite.

BDNF antibody was from Alomone Lab (#ANT-010) or R&D Systems (#MAB648), PGC-1α antibody was from Abcam (#ab106814) or Novus Biologicals (#NBP1-04676), YY1 was from Santa Cruz Biotech (#sc-7341), and CPT1b was from Proteintech (#22170-1-AP). Antibodies from Cell Signaling Technologies: (Erk #4695, p-Erk #4370, CREB #9197, p-CREB #9198, AMPK #5832, p-AMPK #4185, AKT #4691, and p-AKT #4060).

NRCMs were transfected with TrkB and YY1 siRNA (Origene # SR506114 and #SR501936) using Lipofectamine™ RNAiMAX Transfection Reagent (Thermo Fisher Scientifics) according to Manufacture’s protocol. The plasmid containing BDNF cDNA (pCMV6 BDNF, Addgene #39857) was a gift from Dr. Barde^46^. The plasmid with YY1 cDNA (pcDNA3.1 HA-YY1, Addgene # 104395) was a gift from Richard Young^47^. Transfection of BDNF or YY1 plasmid was achieved with Lipofectamine 3000 (Thermo Fisher Scientifics) according to Manufacture’s protocol. Isopreoterenol 10μM, LY294002 20μM (Cell signaling Technologies #9901) and rapamycin 50nM (Selleckchem #S1039) were added to NRCMs 24 hours after transfection of siRNA or plasmid. The duration of isoproterenol, LY294002 and rapamycin incubation was 48 hours.

### Chromatin immunoprecipitation assay

Chromatin immunoprecipitation assay (ChIP assay) was performed using SimpleChIP Plus Enzymatic Chromatin IP kit (CST, Cat#9005) according to the manufacturer’s instruction. Briefly, approximately 2.5×10^6^ neonatal myocytes were harvested and fixed with 1% formaldehyde for 10 min. Chromatin was then digested to mononucleosome using micrococcal nuclease(37 °C 20min), followed by sonication using a Bioruptor (Diagenode) with 5 cycles (30 seconds on followed by 30 seconds off). After centrifugation (10,000 × g, 10 min), the supernatant was used for ChIP assay. Each sample will be aliquoted separately for use as input DNA in quantitative PCR analysis. The remainder of each sample were incubated with either YY1 antibody (Cell Signaling Technologies, #46395), rabbit IgG (negative control) or anti-histone 3 antibody (positive control), and Protein A Dynabeads (Thermo Fisher Scientifics) overnight. After extensive wash, the bound DNA was eluted from the protein A Dynabeads and the crosslink was reversed by the incubation with freshly made elution buffer (1% SDS, 0.1 M NaHCO3) and Proteinase K at 65°C for 4 hours. The DNA was extracted and purified with UltraPure™ Phenol:Chloroform:Isoamyl Alcohol (25:24:1, v/v), and was used as PCR template. Quatitative PCRs were performed with primers listed below: 5’-AAG CGT TAC TTC ACT GAG GCA GAG G-3’ (forward) and 5’-ACG GCA CACACT CAT GCA GGC AAC C-3’ (reverse).

### Quantitative RT-PCR

Total RNA was extracted from either NRCMs or snap-frozen heart tissues using TRIzol reagent (Thermo Fisher Scientifics). Reverse transcription was conducted using High-Capacity cDNA Reverse Transcription Kit (Thermo Fisher Scientifics). Taqman primers (Thermo Fisher Scientifics) were used for quantitative RT-PCR analysis: mouse PPARGC1A (Mm01208835_m1), mouse PPARA (Mm00440939_m1), mouse ESRRA (Mm00433143_m1), and mouse TFAM (Mm00447485_m1).

### Statistics

Data were compared within the groups using one-way or two-way ANOVA with Tukey’s Post Hoc Test using GraphPad Prism version 9.0. Unpaired student’s *t* test was used for comparison between two groups. All tests were two-tailed and a P value of less than 0.05 was considered significant. All values are represented as the mean ± SEM.

## Acknowledgements

This work was supported by the National Institutes of Health grant K08 HL130604, American Heart Association Innovative Project Award #18IPA34170219, UPMC Competitive Research Fund (Dr. Ning Feng), and K08HL157616, Samuel and Emma Winters Foundation (Dr. Manling Zhang).

We thank the University of Pittsburgh Rodent Ultrasonography Core and the funding source for the instrument (NIH 1S10OD023684; Advanced High-Resolution Rodent Ultrasound Imaging System), as well as the Center for Metabolism and Mitochondrial Medicine through support from the Pittsburgh Foundation (MR2020 109502). We thank Dr. Charles McTiernan for providing the human heart tissues. We are grateful for Dr. Mark Anderson and Dr. Toren Finkel’s critical review of the manuscript.

## Author information

### Contributions

M.Z. and N.F. designed the experiments. XY, MZ and NF performed most of the experiments and wrote the manuscript. B.X., M.J. and I.S. conducted isolated working heart experiments and interpreted the data. R.Z., M.K., Z.P and M.O. performed exercise study and some molecular biology experiments. Q.W, J.B. and M.Z. performed echocardiography and analyzed the data. A.W. and B.O. performed mitochondrial physiology study from heart tissues. M.Z. and G.Z. performed TAC surgery. M.R. and S.S. did mitochondrial energetics experiments in cultured cardiomyocytes. H.J., I.S., N.P. and D.A.K. participated in data interpretation and manuscript editing.

## Data availability

The data in support of the findings of this study may be found within the manuscript and in the associated supplementary files. Data associated with this study will be made available from the corresponding author upon reasonable request.

## Supplementary information

**Figure S1.**
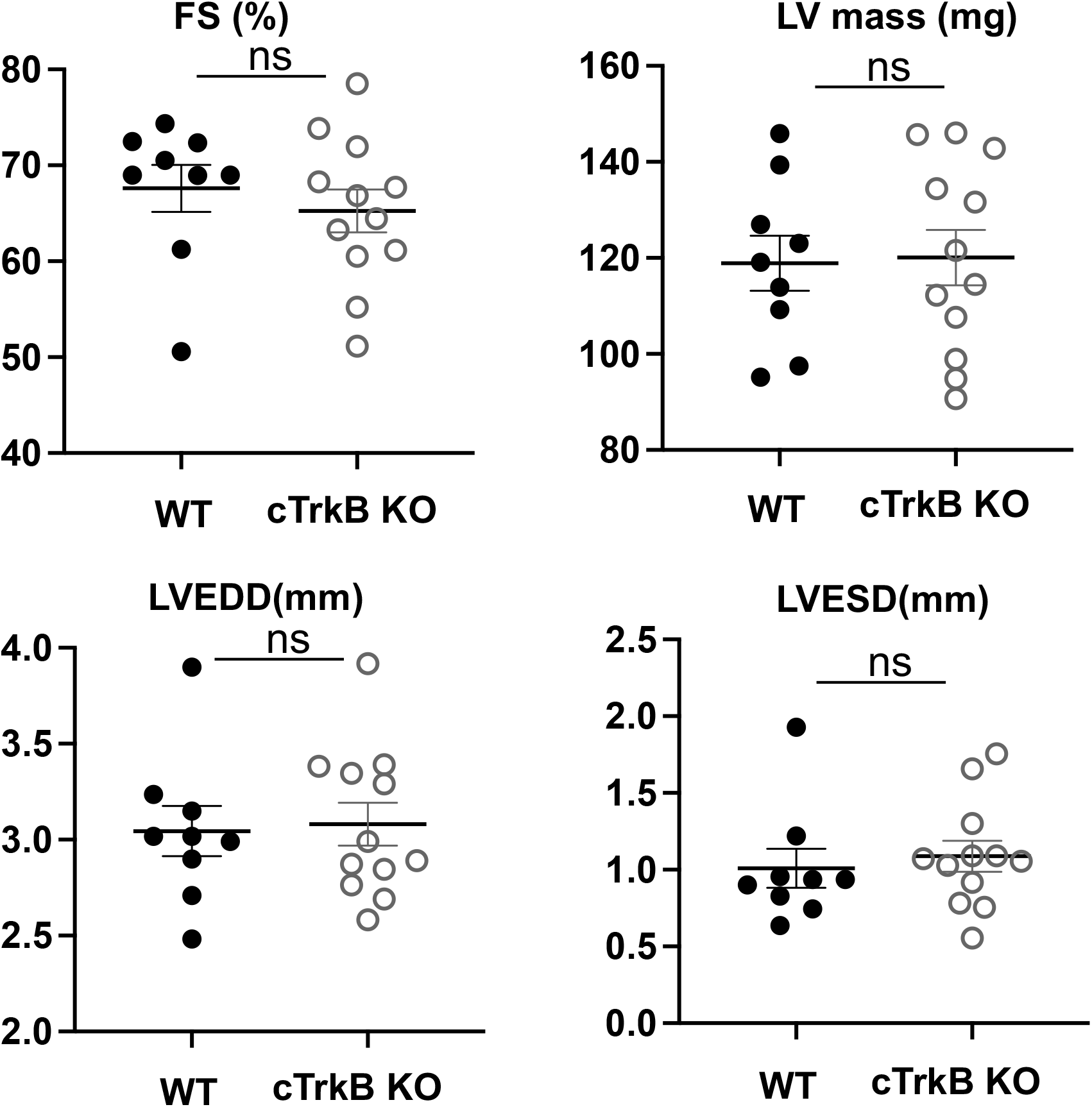
The cardiac function of one-year old cTrkB KO mice is preserved. The fractional shortening, LV mass, left ventricular end systolic or diastolic dimension of cTrkB KO mice were within normal limits. There is no difference in the cardiac function between the WT littermate controls and cTrkB KO mice at the age of one year old (n=9-12).

**Figure S2.**
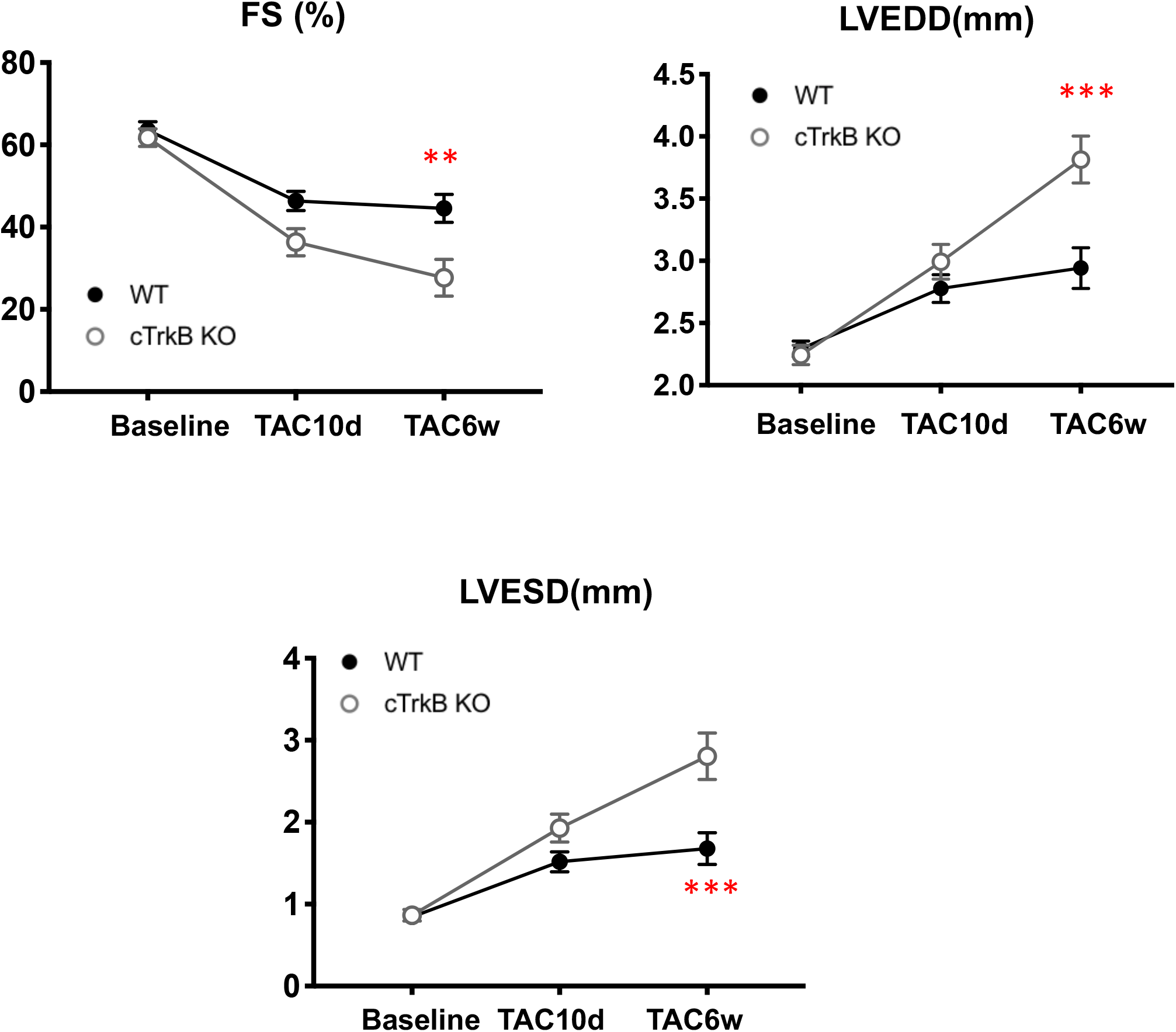
Female cTrkB KO mice show accelerated heart failure progression post-TAC. Compared to WT mice, female cTrkB KO mice showed decreased fractional shortening (WT 44.6±3.4% vs cTrkB KO 27.7±4.4%, P=0.0062), increased LV end diastolic dimension (LVEDD, WT 2.94±0.16 mm vs cTrkB KO 3.8±0.2 mm P=0.0006) and LV end systolic dimension (LVESD, WT 1.7±0.2 mm vs cTrkB KO 2.8±0.3 mm, P=0.0002). (WT n=11, cTrkB KO n=7). *\*\*P<0.01 ***P<0.001*

**Figure S3.**
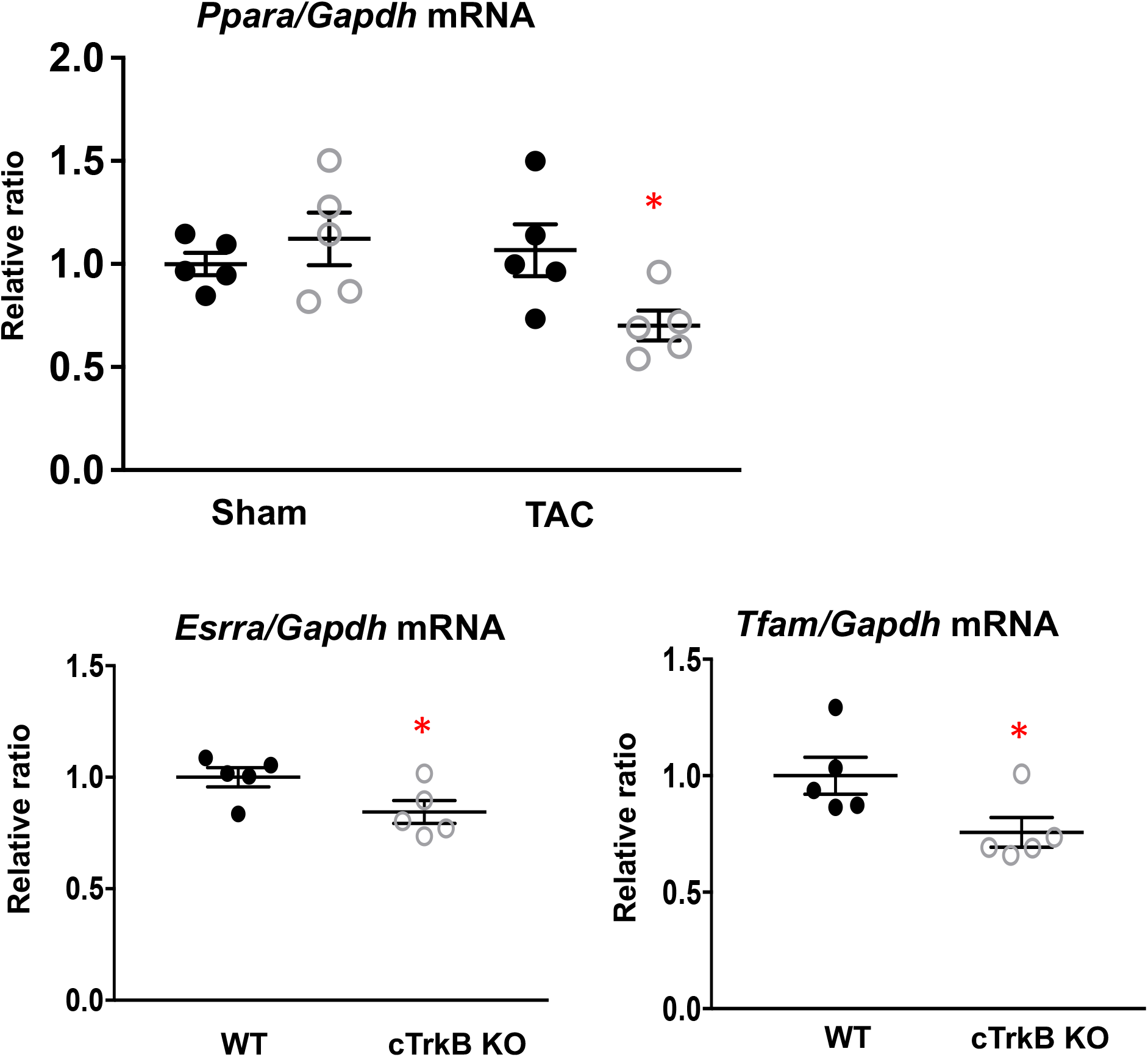
PPARα, ERRα and TFAM expression are further decreased in TAC cTrkB KO mice. Compared to WT controls, metabolic transcription factor PPARα (Ppara), ERRα (Essra) and TFAM mRNA level were further decreased in cTrkB KO mice with TAC. (n=5, *\*P<0.05*)

**Figure S4.**
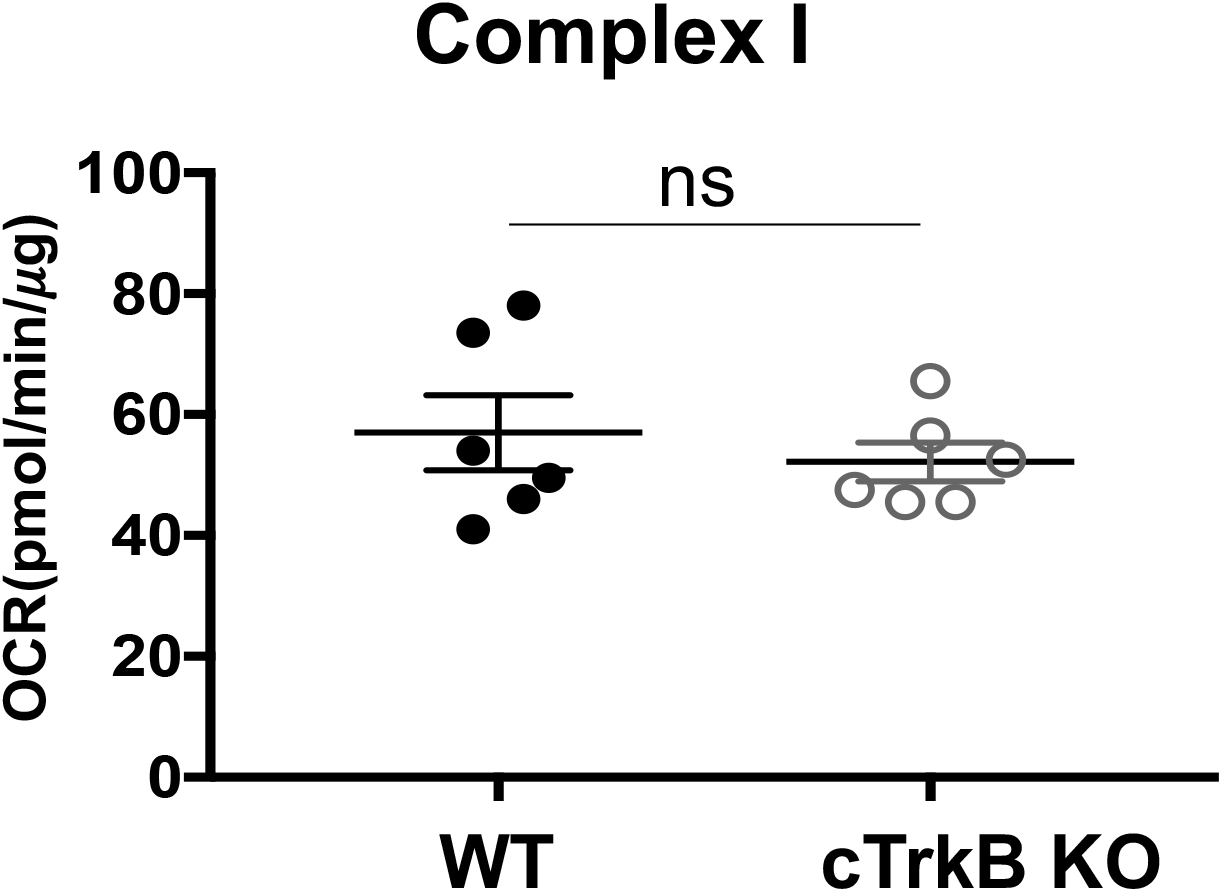
Complex I oxygen consumption rate is preserved in cTrkB KO mice subjected to TAC. There is no difference in complex I respiration between WT and cTrkB KO mice after TAC. (n=6).

**Figure S5.**
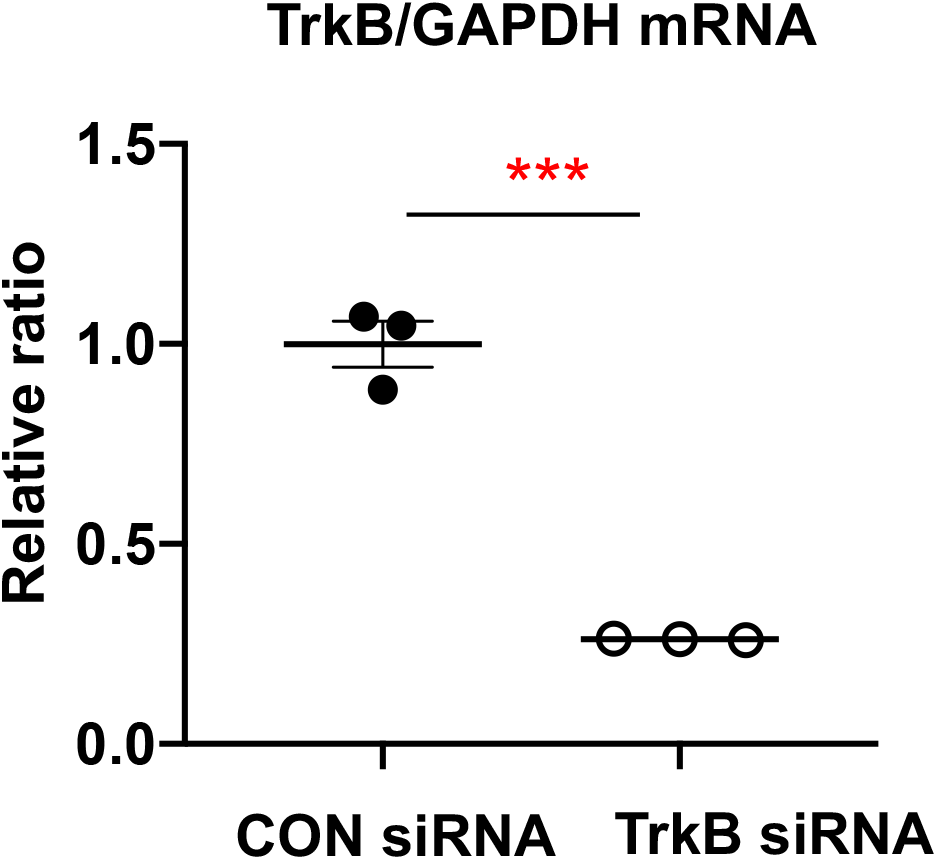
The efficiency of TrkB knockdown with siRNA in NRCMs. Quantitative RT-PCR showed TrkB mRNA was decreased by 75% with siRNA in NRCMs.

**Figure S6.**
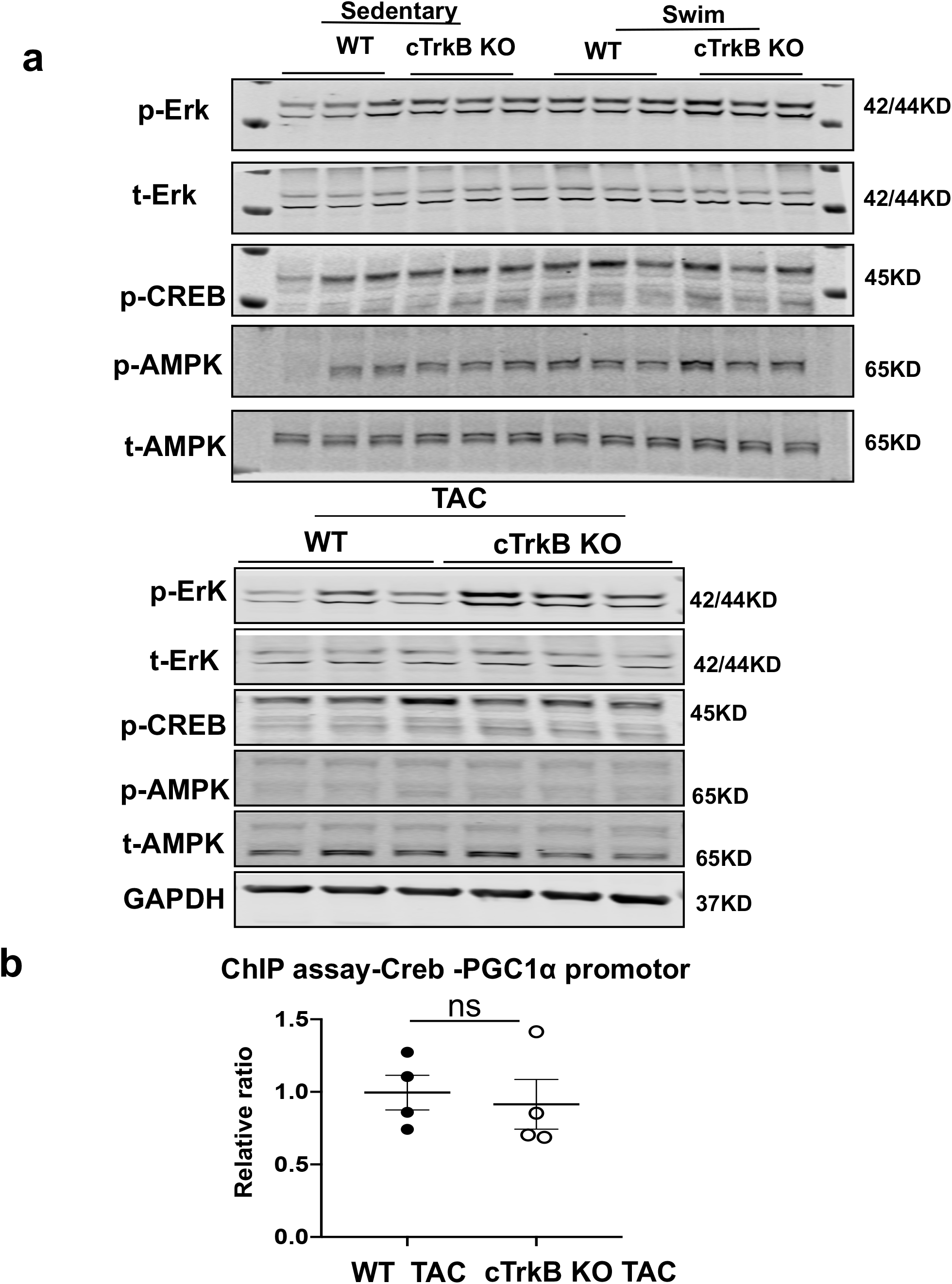
Erk, CREB, and AMPK signaling pathways in WT or cTrkB KO mice subjected to swimming exercise or TAC. **a.** There was no significant difference in the level of phosphorylated or total Erk, CREB, AMPK between WT and cTrkB KO mice subjected to exercise or TAC, except p-Erk was increased in TAC cTrkB KO mice **b.** ChIP assay showed there was no difference in the binding of CREB to PGC-1α promoter between TAC WT and cTrkB KO mice (n=4)

